# MM-Deacon: Multimodal molecular domain embedding analysis via contrastive learning

**DOI:** 10.1101/2021.09.17.460864

**Authors:** Zhihui Guo, Pramod Kumar Sharma, Liang Du, Robin Abraham

**Affiliations:** Microsoft Corporation, Redmond, WA 98052

**Keywords:** molecular embedding, similarity, multimodal, contrastive learning, SMILES, IUPAC, drug similarity, drug-drug interaction

## Abstract

Molecular representation learning plays an essential role in cheminformatics. Recently, language model-based approaches have been popular as an alternative to traditional expert-designed features to encode molecules. However, these approaches only utilize a single modality for representing molecules. Driven by the fact that a given molecule can be described through different modalities such as Simplified Molecular Line Entry System (SMILES), The International Union of Pure and Applied Chemistry (IUPAC), and The IUPAC International Chemical Identifier (InChI), we propose a multimodal molecular embedding generation approach called MM-Deacon (**m**ultimodal **m**olecular **d**omain **e**mbedding **a**nalysis via **con**trastive learning). MM-Deacon is trained using SMILES and IUPAC molecule representations as two different modalities. First, SMILES and IUPAC strings are encoded by using two different transformer-based language models independently, then the contrastive loss is utilized to bring these encoded representations from different modalities closer to each other if they belong to the same molecule, and to push embeddings farther from each other if they belong to different molecules. We evaluate the robustness of our molecule embeddings on molecule clustering, cross-modal molecule search, drug similarity assessment and drug-drug interaction tasks.

## 1 Introduction

Drug discovery process involves screening of millions of compounds in the early stages of drug design, which can be time consuming and expensive. Computer-aided drug discovery can reduce the time and cost involved in this process via automating various cheminformatics tasks [1, 2, 3, 4, 5]. Quantitative representation of molecules, a prerequisite for computer-aided drug discovery [1, 6, 7, 8, 9, 10], embeds molecule as numerical vectors in a high-dimensional space.

Traditional methods for molecule embeddings such as fingerprint generation rely heavily on molecular fragment-level operations [11, 12, 13, 14, 15, 16, 17]. An example of such methods is Morgan fingerprint, also known as Extended-Connectivity Fingerprint (ECFP) [15, 18], where a fixed binary hash function is applied on each atom and its neighborhood. These kind of approaches focus on local features, hence may not capture global information.

Just like the revolution in many other research areas such as image perception [19], speech recognition [20] and natural language processing [21], deep learning [22] has also achieved remarkable success in cheminformatics and drug discovery [23, 24, 25, 26, 27] on a variety of tasks such as adverse drug reaction prediction [26, 28], binding affinity prediction [29, 30] and molecular representation learning [2, 31, 32, 33, 34]. Particularly, advances in natural language processing (NLP), like long short-term memory (LSTM) [35], gated recurrent unit (GRU) [36], variational autoencoder (VAE) [37], and Transformer [38] have been very promising for molecule embedding generation [2, 24, 31, 32, 33, 34, 39, 40, 41, 42].

Xu *et al*. [2] adopted machine translation approach to translate Simplified Molecular Line Entry System(SMILES) strings to itself using encoder-decoder GRUs, where embeddings in the latent space right after the encoder were regarded as sequence to sequence fingerprint. [31, 32] explored latent space of VAE. Samanta *et al*. [31] used VAE to reconstruct SMILES from itself, and Koge *et al*. [32] used VAE to encode similarity distances based on constructed similar and dissimilar molecular pairs in the latent space with metric learning. SMILES Transformer [33] is similar to [2] except that it replaced GRU with Transformers. FragNet [34] also has a similar architecture with SMILES Transformer [33] while it enforced extra supervision to the latent space with augmented SMILES and contrastive learning. It is important to note that all of the machine translation-based methods mentioned above operate in an encoder-decoder setting with SMILES representation as the input to the encoder. Therefore, the underlying chemical knowledge in the embedding is limited to one modality i.e. SMILES representation only. Transformers trained with self-supervised masked language modeling (MLM) loss [38] in chemical domain [43, 44, 45, 46, 47, 48] have also been used for molecule representation learning. However, pretraining objectives like MLM loss tend to impose task-specific bias to the final layers of Transformers [49], which may limit the generalization of the embeddings.

In recent years, contrastive learning has been successful in multimodal vision and language research [50, 51, 52, 53, 54, 55, 56, 57, 58]. Radford *et al*. [50] used image-text pairs to learn scalable visual representations. Carlsson *et al*. [49] showed the superiority of contrastive objectives in acquiring global (not fragment-level) semantic representations.

Inspired by these advances in contrastive learning, we propose MM-Deacon (**m**ultimodal **m**olecular **d**omain **e**mbedding **a**nalysis via **con**trastive learning). Generated embeddings from MM-Deacon capture global context as opposed to fragment level information from traditional methods [50]. Moreover, MM-Deacon utilizes multimodal information, whereas existing deep learning-based molecular embedding methods [2, 31, 32, 33, 34] are limited to single modality only.

MM-Deacon uses Transformers as base encoders to encode SMILES and International Union of Pure and Applied Chemistry (IUPAC) based molecule descriptors and projects embeddings from encoders to a joint embedding space. Then, contrastive learning is used to push the embeddings of positive cross-modal pairs (SMILES and IUPAC from the same molecule) closer to each other and the embeddings of negative cross-modal pairs (SMILES and IUPAC from the different molecules) farther from each other. SMILES is widely used to represent molecule structures as ASCII strings [59, 60] in atom and bond level, while IUPAC nomenclature serves the purpose of systematically naming organic compounds by spoken words that indicate the structure of the compound to facilitate communication [61]. Here instead of using SMILES and IUPAC for sequence-to-sequence translation purpose [62, 63, 64], we obtain positive and negative SMILES-IUPAC pairs and contrast their embeddings at global molecule level instead of fragment level. In this way, different descriptors of molecules are integrated to the same joint embedding space and thus the embeddings are expected to maximize mutual information of SMILES and IUPAC molecule descriptors.

we pretrained MM-Deacon on 10 million molecules randomly selected from PubChem [65] dataset, and used the pretrained MM-Deacon to generate molecular embeddings for downstream tasks. Our main contributions are as follows:

- We propose a novel approach called MM-Deacon for utilizing multiple modalities to generate molecule embeddings using contrastive learning.
- We conduct extensive experiments on multiple tasks: molecule clustering, cross-modal molecule search, drug similarity assessment and drug-drug interaction, and show that our approach outperforms baseline methods and the existing state-of-the-art approaches.

## 2 Materials and Methods

MM-Deacon is a deep neural network designed for SMILES-IUPAC joint learning with an objective to maximize mutual information across modalities, where SMILES and IUPAC depicting the same molecule are enforced to be represented by the same normalized vector in the embedding space, whereas SMILES and IUPAC from different molecules are represented by orthogonal vectors. Thus the similarity among molecules can be measured by pairwise cosine similarity. Transformer encoders with multi-head self-attention layers are utilized to encode SMILES and IUPAC strings. Embeddings from the encoders are pooled globally and projected to the joint embedding space. MM-Deacon is pretrained on 10 million molecules from PubChem.

To demonstrate the effectiveness of our approach, we conduct evaluations on the following tasks:

1. **Molecule clustering**: For molecule clustering, we define five functional groups based on IUPAC names, and analyze the sensitivity of the embeddings towards these functional groups via clustering.
2. **Cross-modal molecule search**: cross-modal (SMILES-to-IUPAC, IUPAC-to-SMILES) search is performed on a test set of 100K molecules selected from PubChem (mutually exclusive from the training set).
3. **Drug similarity assessment**: we provide in-depth analysis of our molecular embeddings obtained from MM-Deacon on drug similarity assessment task on FDA approved drug list, focusing on two candidate drugs and their potential alternatives.
4. **Drug-drug interaction prediction**: The goal of this task is to predict if there exists an interaction between two candidate drugs. MM-Deacon embeddings are used as input drug descriptors.

An overview of our training and evaluation scheme is presented in Figure 1. RDKit [66] is used to generate 2D structure visualization of molecules, for the purpose of interpretability.

**Figure 1:**
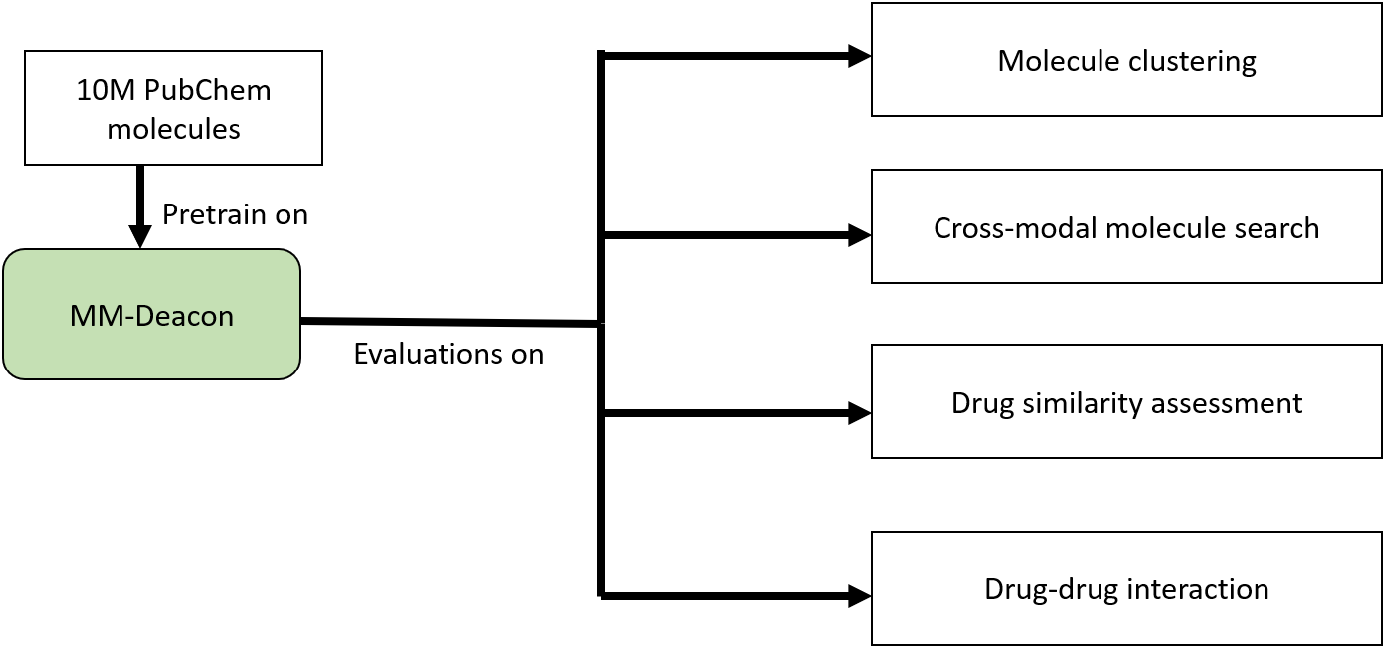
MM-Deacon training and evaluation scheme.

### 2.1 Datasets

There are three datasets used in this study: PubChem, an FDA-approved drug list [31, 34, 67, 68, 69], and a drug-drug interaction dataset [70]. PubChem is a large-scale publicly available dataset that contains the information about millions of chemical compounds and their activities [65]. Molecules that have both canonical SMILES and preferred IUPAC name in their descriptors were extracted, which resulted in 100M SMILES-IUPAC pairs. 10M/100K/100K pairs from the 100M were randomly selected for training/validation/test respectively.

The FDA-approved drug list used here contains 1497 small molecule drugs that were approved before Nov 2013 [31, 34, 67, 68, 69]. Similar to prior studies [31, 34], we also select two drugs as study candidates: an antipsychotic agent called Clozapine [71] and an antibiotic called Flucloxacillin [72]. In this study, we look for their relationships with their potent transporter inhibitors and alternatives in the joint embedding space. For Clozapine, a first-generation typical antipsychotic similar to Clozapine (Loxapine) [71], several drug transporter inhibitors that reduce the uptake of Clozapine (Olanzapine, Quetiapine, Prazosin, Lamotrigine) [31, 73], and a few drugs that have similar structure with Clozapine (Prochlorperazine, Clomipramine, Mirtazapine) are considered as the drugs of interest. For Flucloxacillin, several Penicillin antibiotics (Dicloxacillin, Cloxacillin, Oxacillin, Amoxicillin), a Tetracycline antibiotic treating bacterial infections (Doxycycline) [74], and an appropriate alternative of Flucloxacillin for common skin infections (Erythromycin) [75] are highlighted for comparison.

The drug-drug interaction dataset [70] contains 548 drugs, 48,584 known interactions, and 101,294 non-interactions (may contain undetected interactions). Along with the drug-drug interaction matrix, there are also eight types of feature based drug-drug similarity matrices, which are similarities based on substructure, target, enzyme, transporter, pathway, indication, and off side effect [70]. The substructure similarity matrix is generated from 881 dimensional substructure vectors extracted from PubChem.

### 2.2 MM-Deacon

MM-Deacon takes SMILES and IUPAC strings as the input, where the strings are first split into lists of tokens by dedicated tokenizers. Afterwards, SMILES and IUPAC token vectors are encoded individually by separate Transformers, and then both modalities are projected to a joint embedding space. If SMILES and IUPAC features belong to the same molecule they are identified as positive pair, otherwise they are considered as negative pairs. Contrastive loss is enforced to bring shared features of positive SMILES-IUPAC pairs closer and push negative SMILES-IUPAC pairs farther in the joint embeddings space. The model diagram is shown in Figure 2.

**Figure 2:**
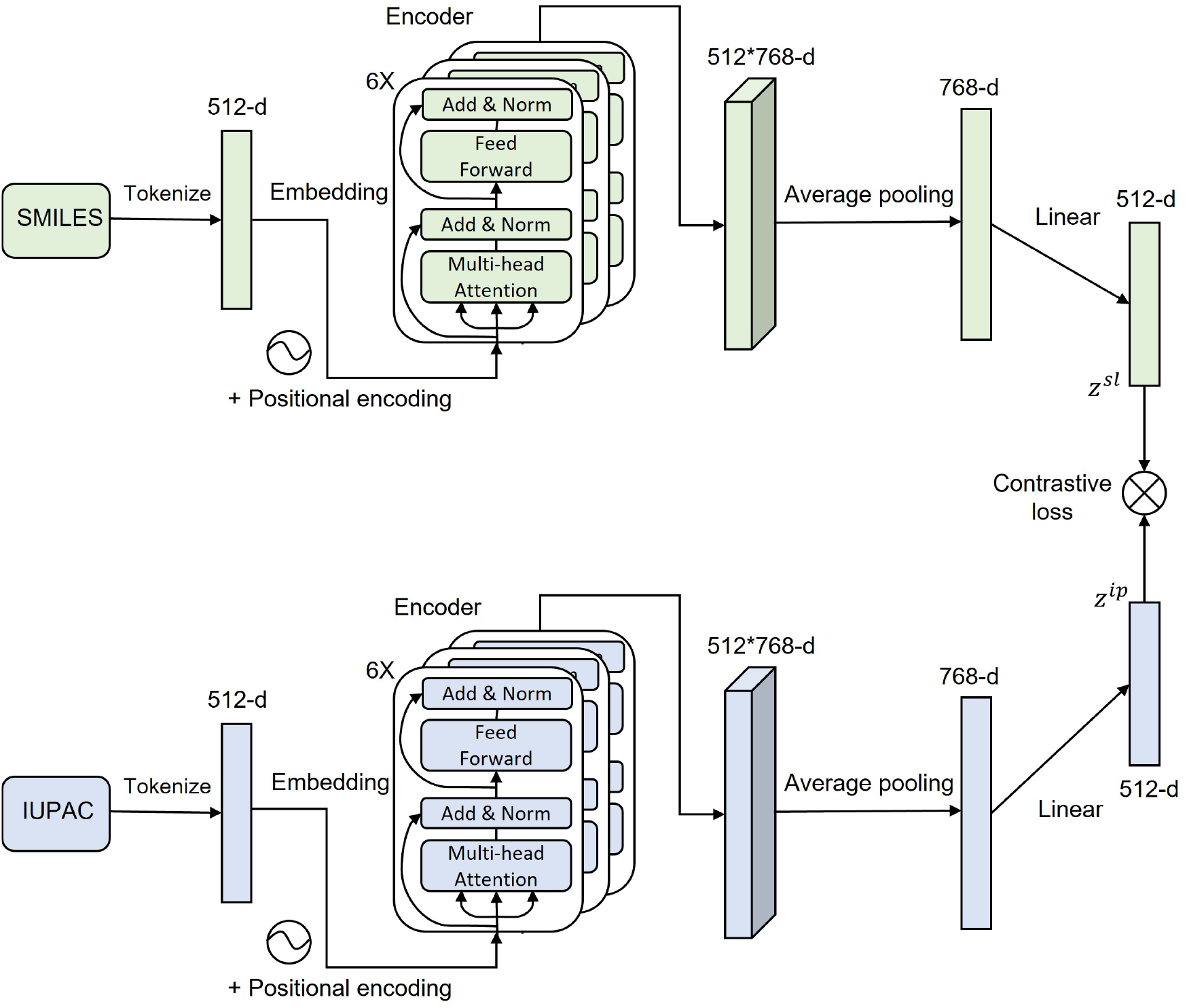
Diagram of MM-Deacon. Green blocks are for SMILES branch and blue blocks are for IUPAC branch. 512-d means 512 dimensions and 6X represents 6 times. *z^sl^* and *z^ip^* means SMILES embedding and IUPAC embedding respectively.

#### 2.2.1 Tokenizers

For SMILES tokenization, we use a Byte-Pair Encoder (BPE) [76, 77] as used in [44], where the authors showed that BPE performed better than regex-based tokenization for SMILES on a downstream task. For IUPAC name tokenization, we use a rule-based regex that splits IUPAC strings according to their suffixes, prefixes, trivial names etc as designed in [63].

#### 2.2.2 Model architecture

As shown in Figure 2, MM-Deacon takes SMILES and IUPAC strings as the inputs to separate branches. Within each branch, the input text string *s* is tokenized and embedded into a numeric matrix representation *x*, and the order of the token list is preserved by a positional embedding *p_x_*. Then *x* and *p_x_* are ingested by an encoder that consists of 6 Transformer encoder blocks. A Transformer encoder block has two sub-layers, a multi-head attention layer and a fully-connected feed-forward layer. Each sub-layer is followed by a residual connection and layer normalization to normalize input values for all neurons in the same layer [38, 78]. The multi-head attention layer acquires long-dependency information by taking all positions into consideration. We then use a global average pooling layer to integrate features at all positions and a linear layer to project the integrated feature vector to the joint embedding space. Thus the final embedding *z* of *x* can be expressed as

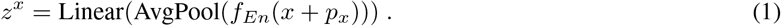

The maximum length of the input token sequence size is set as 512. We choose the number of self-attention heads as 12 and hidden size of 768 for each Transformer encoder block. The final linear layer projects the vector from length of 768 to 512 to make the representation more compact. Thus *z^x^* ∈ ℝ^512^.

#### 2.2.3 Contrastive loss

Our objective is to align pairs of modalities by maximizing mutual information of positive pairs, and discriminating them from negative pairs in the joint embedding space. For this purpose, we use InfoNCE [50, 55, 79] as the contrastive loss. We do not construct negative pairs manually. Instead, we obtain negative pairs in minibatches during training. Considering a minibatch of *N* SMILES-IUPAC pairs as input, within the correlation matrix of *N* SMILES strings and *N* IUPAC strings, *N* positive pairs and *N*^2^ − *N* negative pairs can be generated. For *i*-th SMILES, the only positive pair is *i*-th IUPAC while the rest *N* − 1 IUPAC strings are all negative pairs. Therefore, the InfoNCE loss for *i*-th SMILES is,

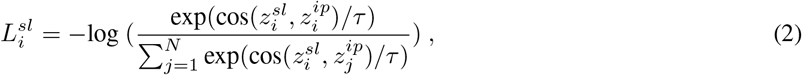

where *sl, ip* represent SMILES and IUPAC respectively. cos() is the cosine similarity function, and *τ* is the temperature. Likewise, the loss function for *i*-th IUPAC is,

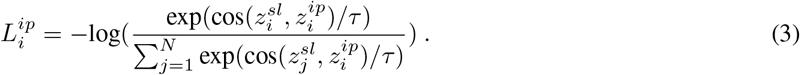

Thus, the final loss function can be written as,

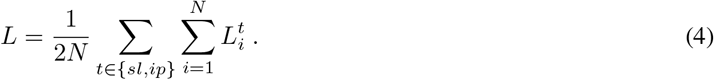

We use AdamW optimizer with a learning rate of 10^−6^, and train MM-Deacon on 80 V100 GPUs for 10 epochs (15 hours) with a 16 batch size on each GPU. The temperature *τ* is set as 0.07 as in [79].

### 2.3 MM-Deacon joint embedding space analysis

After MM-Deacon pretraining has been completed, the joint embedding space is evaluated on four different tasks, as shown in Figure 1. First, clustering analysis in the joint embedding space is provided to check the sensitivity of generated embeddings to domain knowledge in functional group level. For this purpose, we construct five functional groups (nitro, fluoro, chloro, bromo, and sulfonamide) of molecules from the PubChem test set, with the rule that if the IUPAC name of the molecule includes the name of the group and excludes names of all other groups, then the molecule is considered belonging to this group. Characteristics of each group are highlighted in Figure 3.

**Figure 3:**
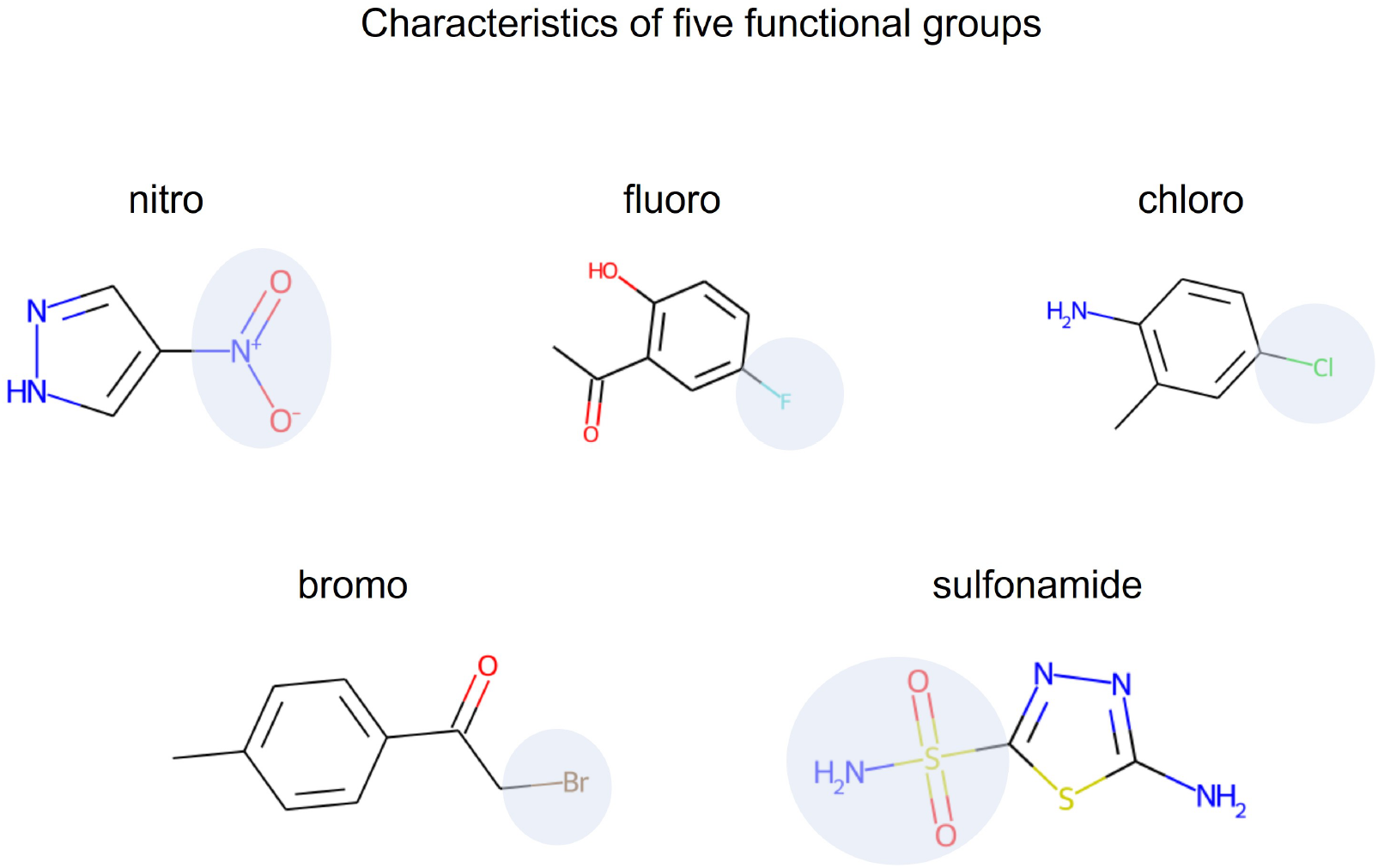
Examples of five distinctive functional groups indicated by their group names. The characteristics of each group are highlighted by light blue circles overlaid on RDKit plots of the sample molecules.

PubChem cross-modal molecule search serves as a way to test the learned agreement across SMILES and IUPAC representations in the joint embedding space. Specifically, molecules in the PubChem test set are all embedded into 512-dimensional vectors in the joint embedding space. For a given query vector, cosine similarity scores between the query and search candidates are calculated to determine the ranking. Feature reduction tools like t-SNE [80] and UMAP [81] are used in clustering analysis and cross-modal molecule search to visualize molecular embeddings in 2D plane.

For drug similarity assessment on FDA-approved drug list, to find the similar drugs as query drug candidates, we obtain the molecule embeddings through drug SMILES representation (*z^sl^*) from pretrained MM-Deacon and the drug-drug relationships are mapped via cosine similarity.

Lastly, a drug-drug interaction prediction task is conducted to validate the effectiveness of MM-Deacon in predicting molecular property related tasks. We concatenate SMILES embeddings of the pairwise drugs and use a multi-layer perceptron (MLP) network [82] implemented by scikit-learn [83] to predict the binary labels. The MLP has one hidden layer with 200 neurons. ReLU activation and a learning rate of 10^−3^ are used. Stratified 5-fold cross-validation that balances distribution of number of interactions and non-interactions in each fold is employed to report the final results.

## 3 Results

### 3.1 Molecule clustering analysis

Clustering results based on t-SNE feature reduction of SMILES embeddings and IUPAC embeddings are displayed in Figures 4 and 5 respectively, where molecules in five different functional groups are marked by different colors. For clarity, only 1000 randomly selected data points in each group are selected for clustering. From Figures 4 and 5, it is clear that both SMILES and IUPAC embeddings show excellent separation abilities of molecules at the functional group level. For quantitative evaluation of clustering, we calculated four indices implemented by scikit-learn that measure clustering qualities from different aspects, namely homogeneity, completeness, adjusted mutual information (AMI), and Fowlkes-Mallows index (FMI). Annotated group classes are used as the true labels and predicted labels are generated by k-means clustering (k=5) [84]. Homogeneity measures if one cluster only contains molecules of a single group, whereas Completeness checks if all molecules of the same group are assigned to the same cluster. AMI is the adjusted measurement of the agreement between clustering and true labels, and FMI calculates the geometric mean of the pairwise precision and recall. The higher the indices are, the better the performance is. From Table 1, we can see that SMILES embeddings perform slightly better than the IUPAC embeddings, hence have better separation among the points from these five functional groups.

**Figure 4:**
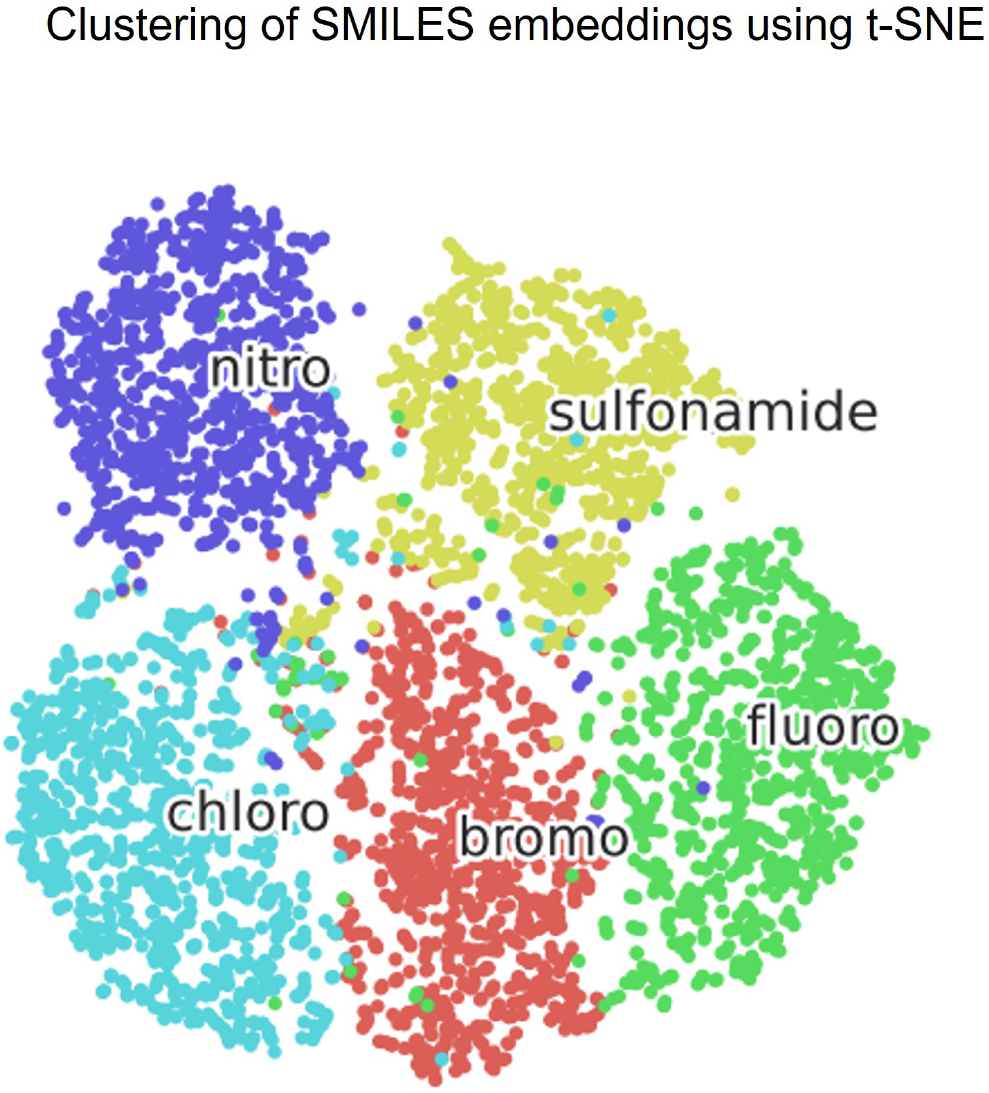
Clustering of SMILES embeddings for molecules in five functional groups using t-SNE. Different groups are represented by different colors.

**Figure 5:**
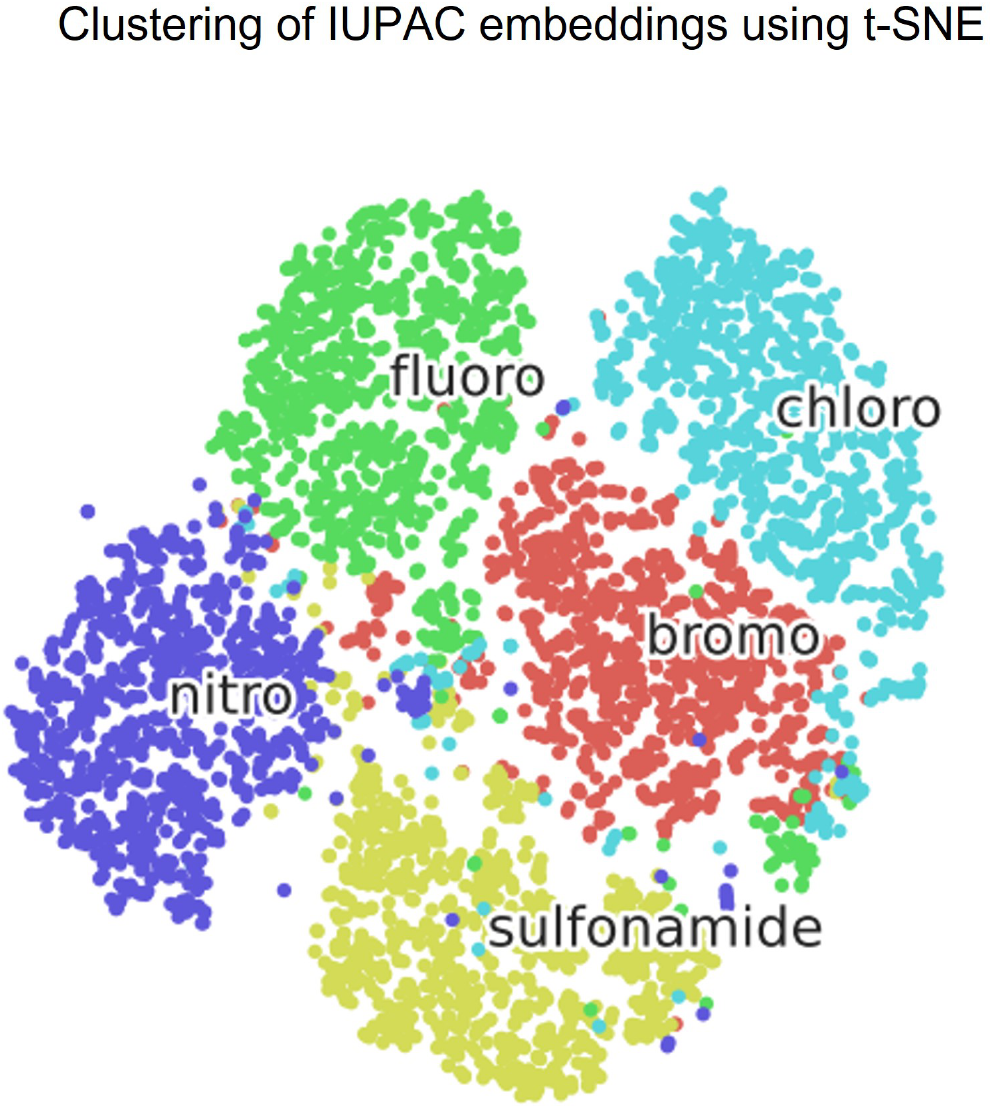
Clustering of IUPAC embeddings for molecules in five functional groups using t-SNE. Different groups are represented by different colors.

**Table 1:** Clustering performance evaluation for different embedding types in the joint embedding space on separating five functional groups.

| Embedding type | Homogeneity | Completeness | AMI | FMI |
| --- | --- | --- | --- | --- |
| SMILES | **0.815** | **0.816** | **0.815** | **0.878** |
| IUPAC | 0.757 | 0.759 | 0.758 | 0.818 |

### 3.2 Cross-modal molecule search

For cross-modal molecule search, we created search corpus in three different settings to give a comprehensive evaluation of the performance. In addition to running search directly on entire Pubchem test set of 100K SMILES-IUPAC pairs, we also reported the search performance averaged on smaller subsets randomly split from PubChem test set, i.e. 1K 100-sized subsets and 10 10K-sized subsets. We used Average Recall at K as the evaluation metric. Average Recall@K measures the percentage of the ground truth appearing in the top K retrieved molecules. For our experiments, we used two values of K(1 and 5). For baseline comparison, besides pretraining on 10M molecules, we also pretrained MM-Deacon on 1M molecules, which is a subset of the 10M training set. An illustration of cross-modal IUPAC-SMILES search is shown in Figure 6. The evaluation results for SMILES-to-IUPAC and IUPAC-to-SMILES search on all three test corpus are shown in Table 2. We can see from Table 2 that MM-Deacon pretrained on 10M molecules performs significantly better as compared to pretraining on 1M molecules.

**Figure 6:**
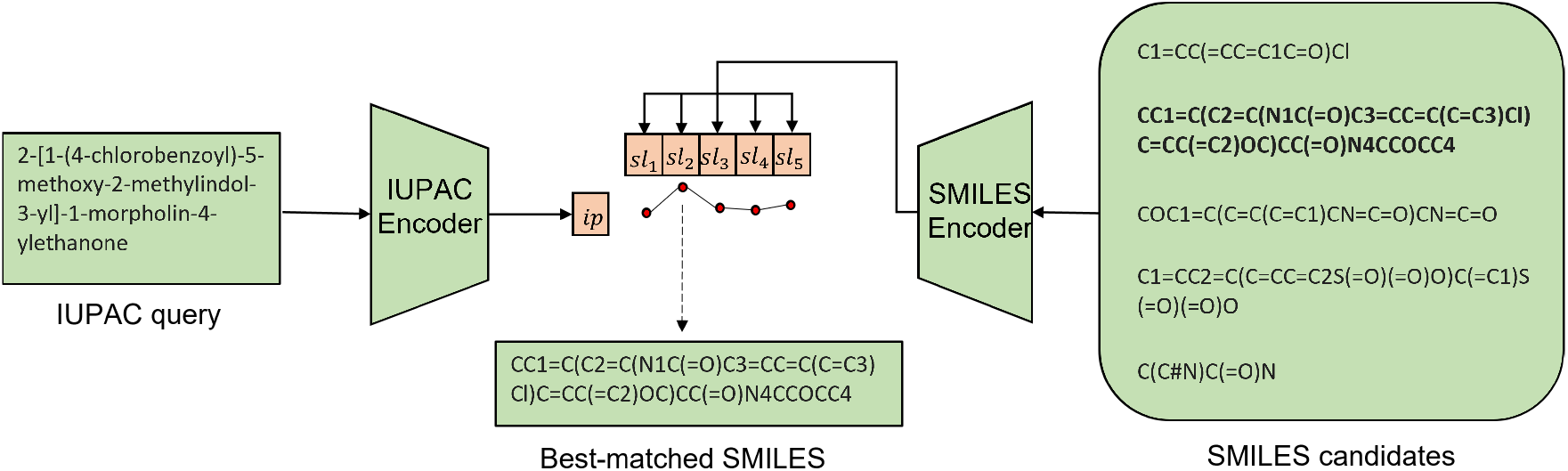
An illustration of zero-shot molecule IUPAC-to-SMILES search. Bold textual candidate is the SMILES ground truth for the query IUPAC. Red dots show the cosine similarity scores between query IUPAC embedding and candidate SMILES embeddings.

**Table 2:** Average recall for cross-modal retrieval on PubChem test set with different search size settings.

|  |  | SMILES-to-IUPAC |  |  | IUPAC-to-SMILES |  |  |
| --- | --- | --- | --- | --- | --- | --- | --- |
|  |  | 100 | 10K | 100K | 100 | 10K | 100K |
| Pretrained on 1M molecules | Recall@1 | 99.67% | 41.83% | 15.97% | 99.89% | 43.26% | 20.01% |
|  | Recall@5 | 100% | 79.42% | 39.80% | 100% | 81.35% | 47.68% |
| Pretrained on 10M molecules | Recall@1 | 99.98% | 84.62% | 52.09% | 100% | 87.46% | 59.92% |
|  | Recall@5 | 100% | 98.97% | 85.36% | 100% | 99.10% | 88.69% |

One example for each of SMILES-to-IUPAC and IUPAC-to-SMILES search on the entire PubChem test set of size 100K is displayed in Figures 7 and 8 respectively. Depending on the search objective, in each figure, either SMILES or IUPAC strings of the top 5 most similar molecules to the query are listed in the panel on the right, and the corresponding embeddings are projected to 2D plane using UMAP in the left panel. Note that for clarity, only top 1K molecules are plotted in the UMAP panel. RDKit plots of each molecule are shown in the table for graphical interpretation. An observation is that critical substructures in the queries are all present in the top 5 retrieved results. Moreover, the top-ranked molecules are very close to each other in the UMAP 2D coordinates.

**Figure 7:**
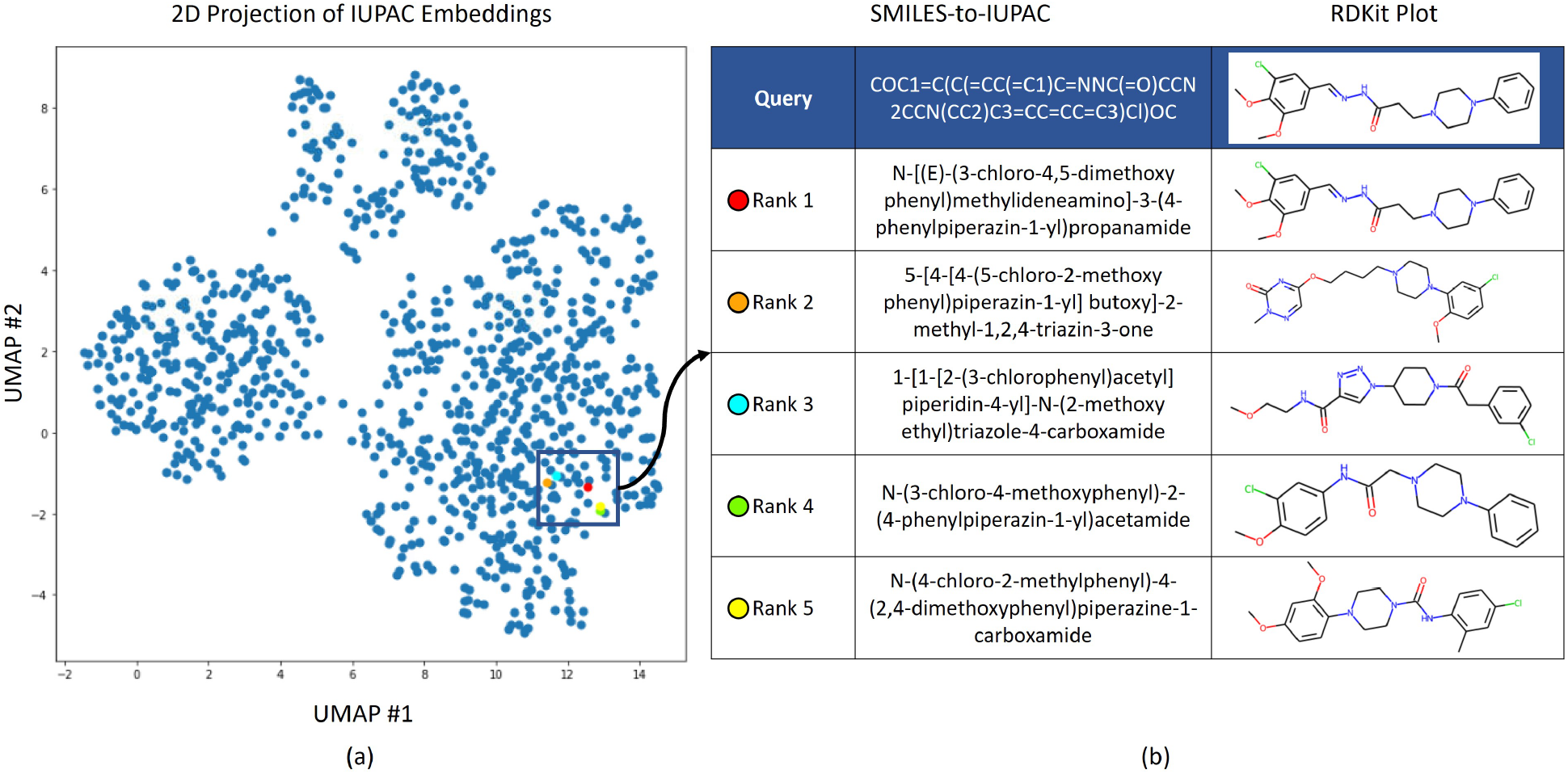
Example for SMILES-to-IUPAC search on 100K molecules. (**a**) 2D projection of IUPAC embeddings using UMAP for top 1K ranked molecules. (**b**) Table with the example SMILES query and top 5 ranked IUPAC names. RDKit plots of corresponding molecules are placed next to each string to facilitate interpretation. The locations of the top 5 molecules in (**b**) are within the black square in (**a**) and marked by different colors.

**Figure 8:**
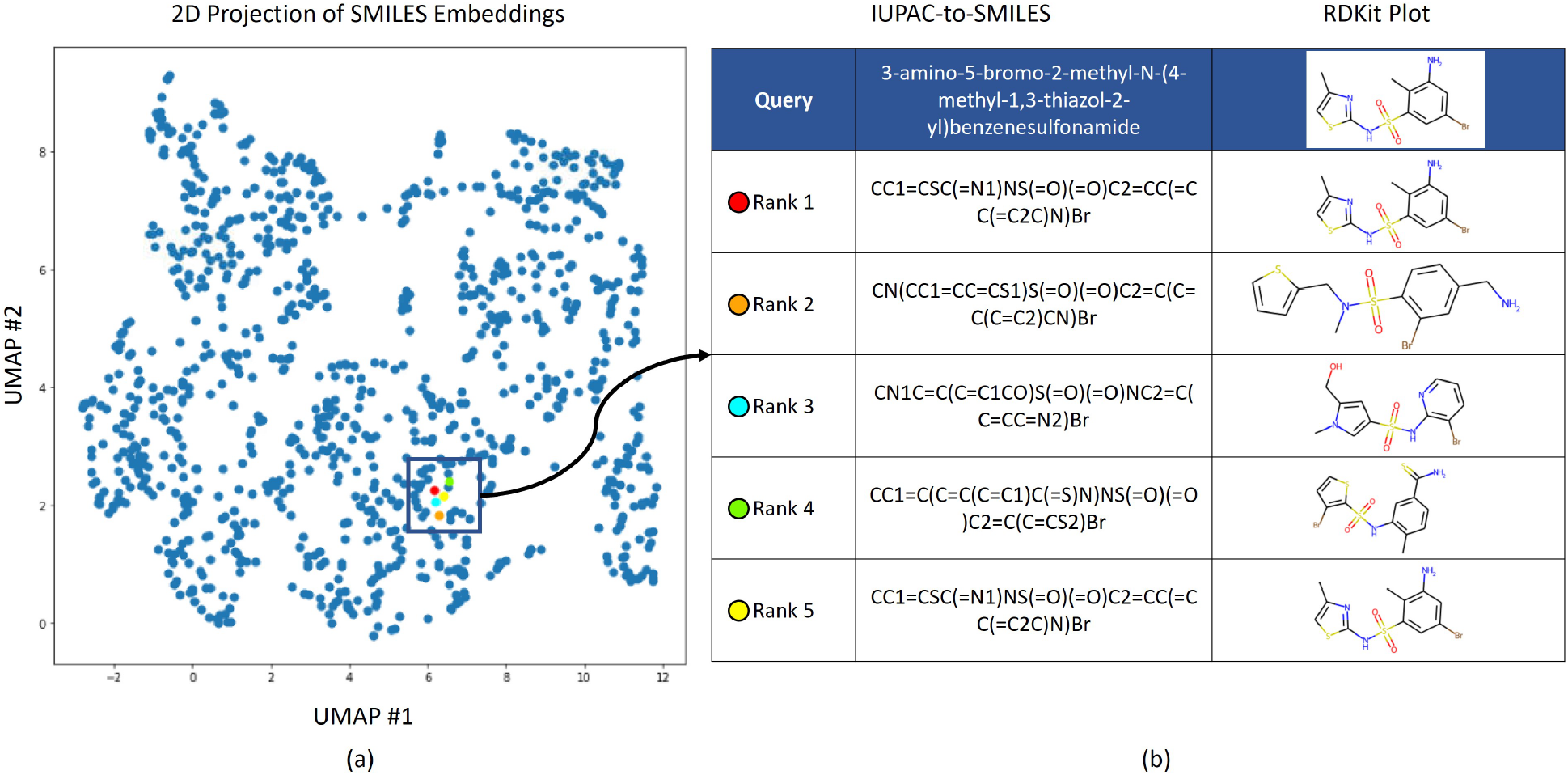
Example for IUPAC-to-SMILES search on 100K molecules. (**a**) 2D projection of SMILES embeddings using UMAP for top 1K ranked molecules. (**b**) Table with the example IUPAC query and top 5 ranked SMILES strings. RDKit plots of corresponding molecules are placed next to each string to help with the interpretation. The locations of the top 5 molecules in (**b**) are within the black square in (**a**) and marked by different colors.

### 3.3 Drug similarity assessment

For drug similarity assessment, we used an FDA approved drug list. First, embeddings are generated in the joint embedding space using MM-Deacon based on the SMILES representations of the drugs. Then cosine distances are computed between the candidate drugs and the other drugs in the dataset. In our experiments, we selected Clozapine and Flucloxacillin as two candidate drugs and analyzed their similarity scores against the drugs of interest mentioned in section 2.1. We compared cosine similarity results with two popular molecular similarity measurements, namely Tanimoto Similarity [85] with RDKit fingerprint and Morgan fingerprint [31].

Similarity measurement from Tanimoto Similarity (RDKit) against Tanimoto Similarity (Morgan) is presented in Figure 9. The similarity scores from both approaches decrease very fast, and both Lamotrigine and Prazosin are low-ranked with scores less than 0.4. Cosine similarity against Tanimoto Similarity (RDKit) is shown in Figure 10 and cosine similariy against Tanimoto Similarity (Morgan) is displayed in Figure 11. From Figures 10 and 11, we can see that Tanimoto Similarity has low ranks for some of the drugs of interest, whereas most of the drugs of interest have a very high rank using cosine similarity.

**Figure 9:**
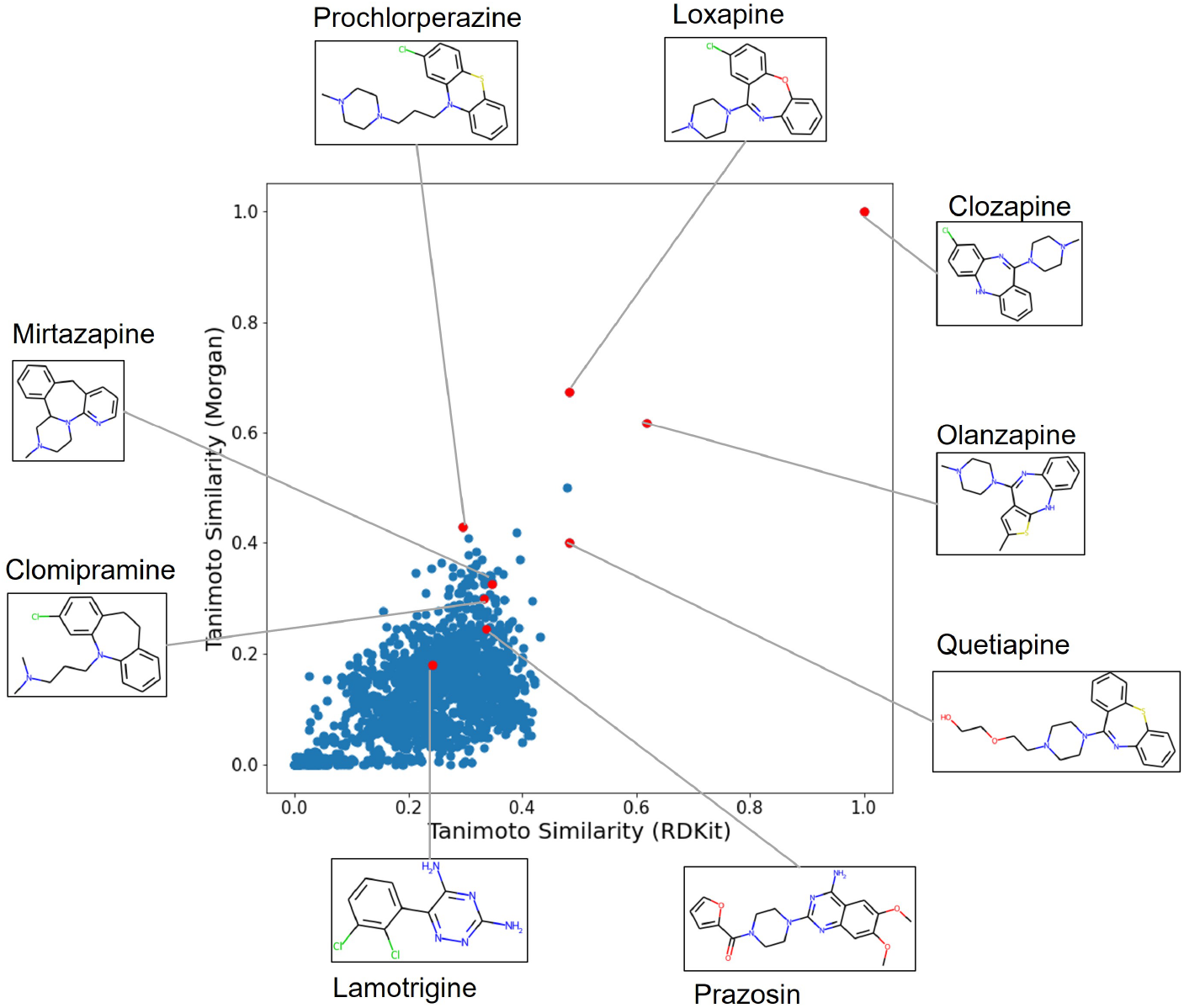
Scatter plot of two kinds of Tanimoto Similarity scores using RDKit fingerprint (x-axis) and Morgan fingerprint (y-axis) between Clozapine and 1497 FDA-approved drugs. Red dots indicate drugs of interest, while blue ones are the rest. The corresponding drug names and RDKit plot of the drug molecules are also shown.

**Figure 10:**
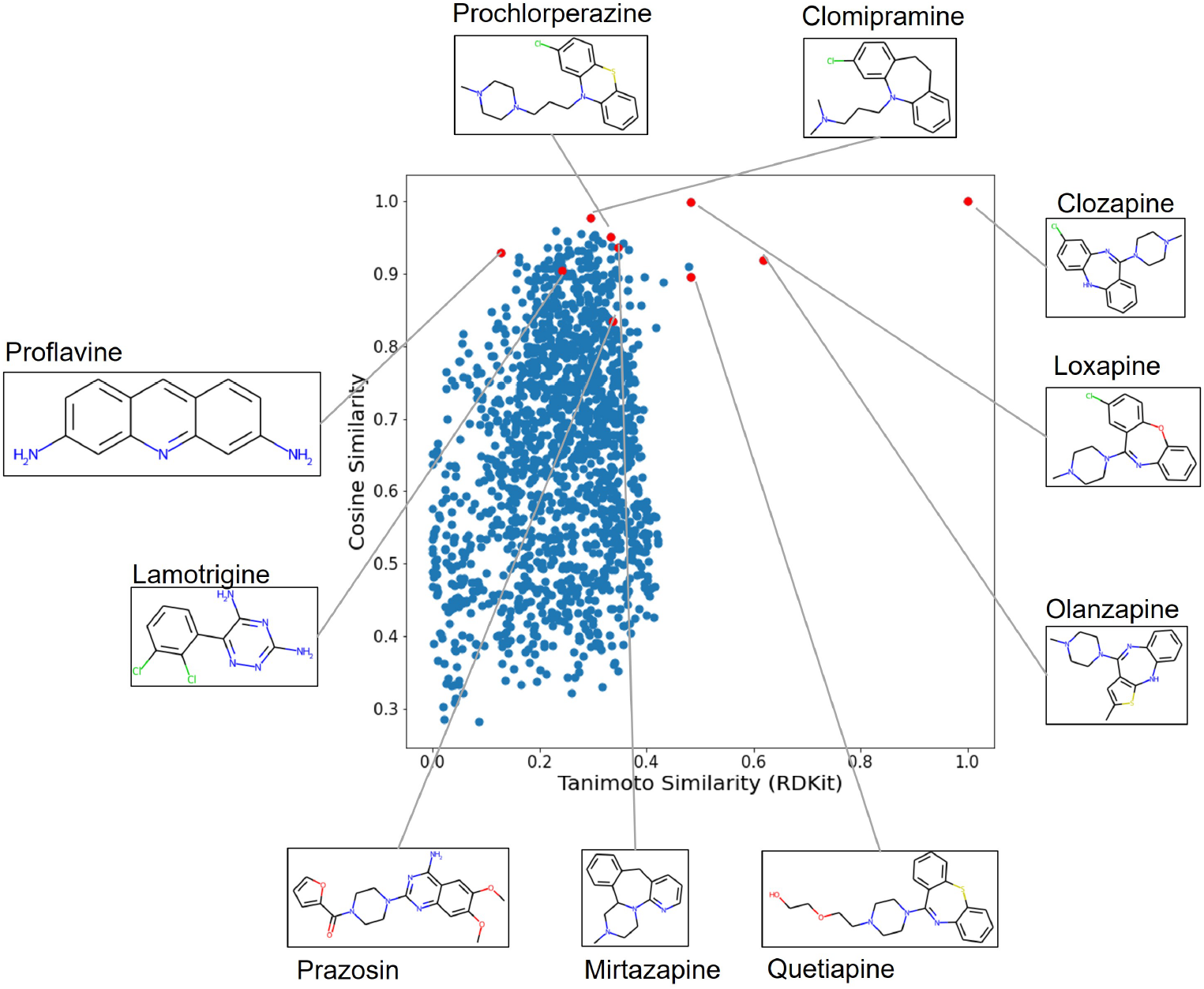
Scatter plot of cosine similarity against Tanimoto Similarity with RDKit fingerprint between Clozapine and 1497 FDA-approved drugs.

**Figure 11:**
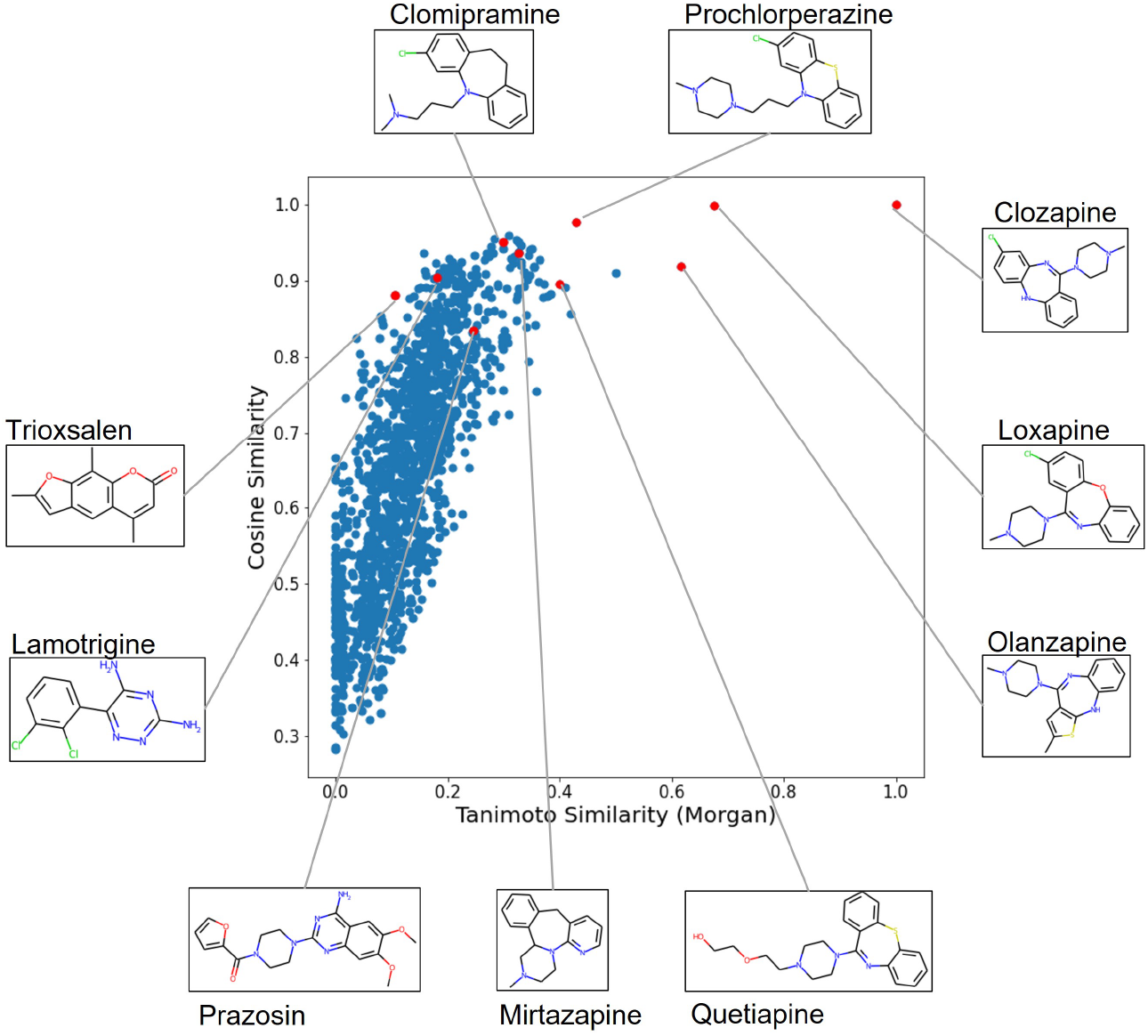
Scatter plot of cosine similarity against Tanimoto Similarity with Morgan fingerprint between Clozapine and 1497 FDA-approved drugs.

In Figure 12, cosine similarity is compared with Euclidean distance. The Euclidean distance here measures the shortest distance between Clozapine embedding with the embeddings of all drugs in a 512 dimensional space, whereas cosine similarity calculates the cosine of the angle between two embeddings. From this figure, it is clear that there is a good alignment of the two different ways to measure closeness in the same space.

**Figure 12:**
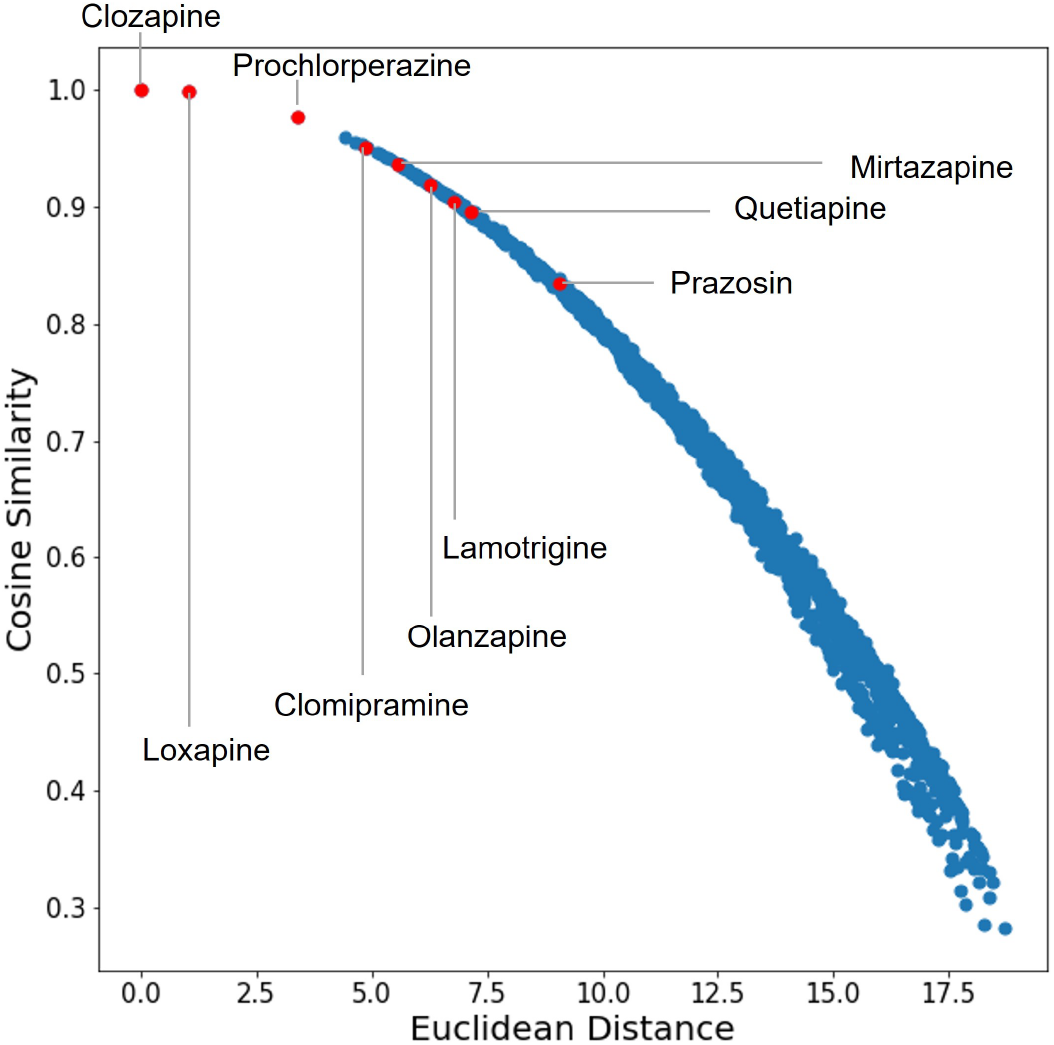
Cosine similarity vs. Euclidean distance in the embedding space between Clozapine and 1497 FDA-approved drugs.

Similarly for Flucloxacillin, pairwise comparisons among cosine similarity, Tanimoto Similarity (RDKit), and Tanimoto Similarity (Morgan) are shown in Figures 13, 14 and 15. From Figures 13 and 15, we can see that a massive number of drugs are in a plateau using Tanimoto Similarity (Morgan) measurement, and Erythromycin has an extremely low rank. The rankings of drugs of interest are comparative for cosine similarity and Tanimoto Similarity (RDKit) as shown in Figure 14. Comparison between cosine similarity and euclidean distance in the embedding space is shown in Figure 16.

**Figure 13:**
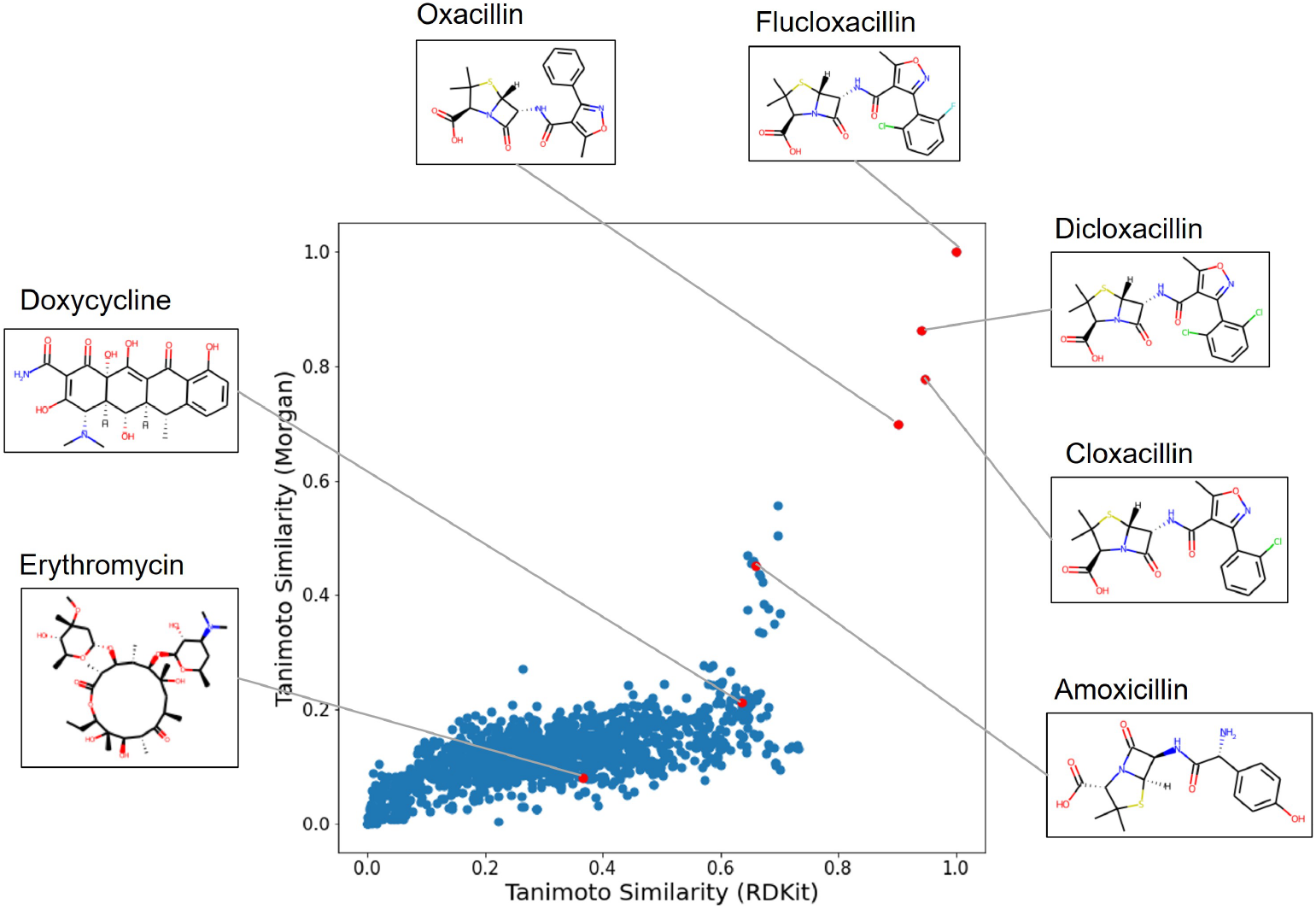
Scatter plot of Tanimoto Similarity with RDKit fingerprint and Morgan fingerprint between Flucloxacillin and 1497 FDA-approved drugs.

**Figure 14:**
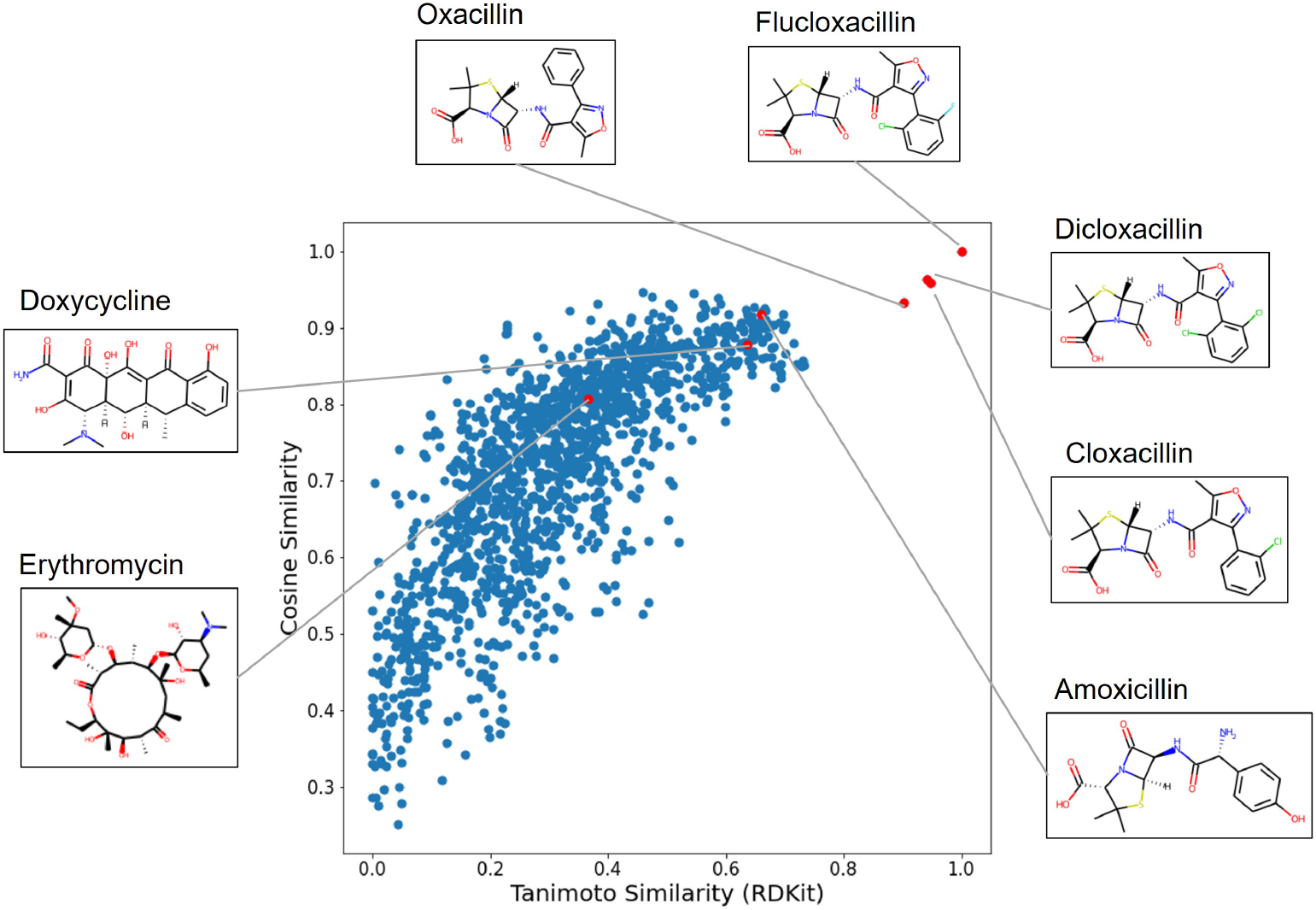
Scatter plot of cosine similarity against Tanimoto Similarity with RDKit fingerprint between Flucloxacillin and 1497 FDA-approved drugs.

**Figure 15:**
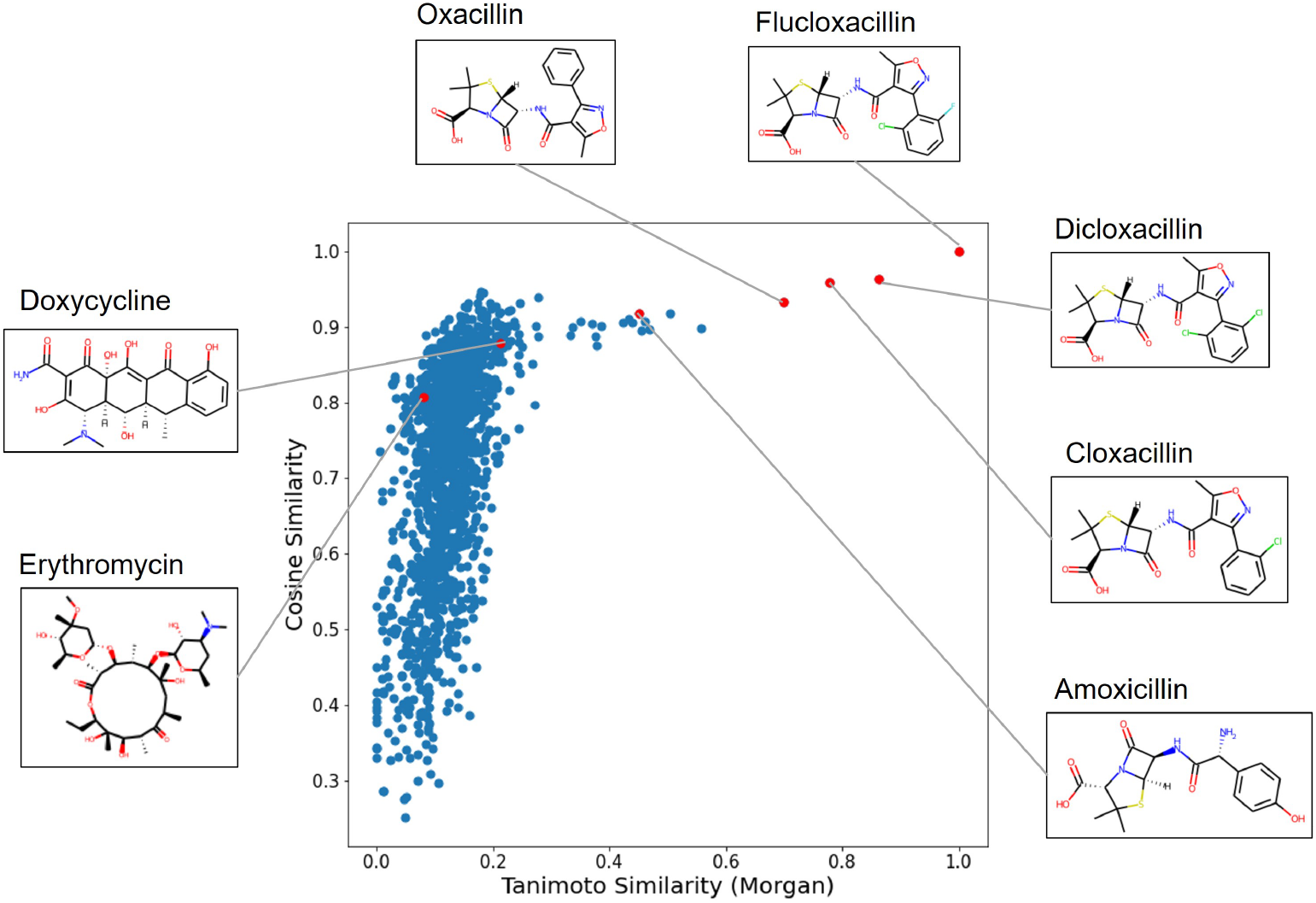
Scatter plot of cosine similarity against Tanimoto Similarity with Morgan fingerprint between Flucloxacillin and 1497 FDA-approved drugs.

**Figure 16:**
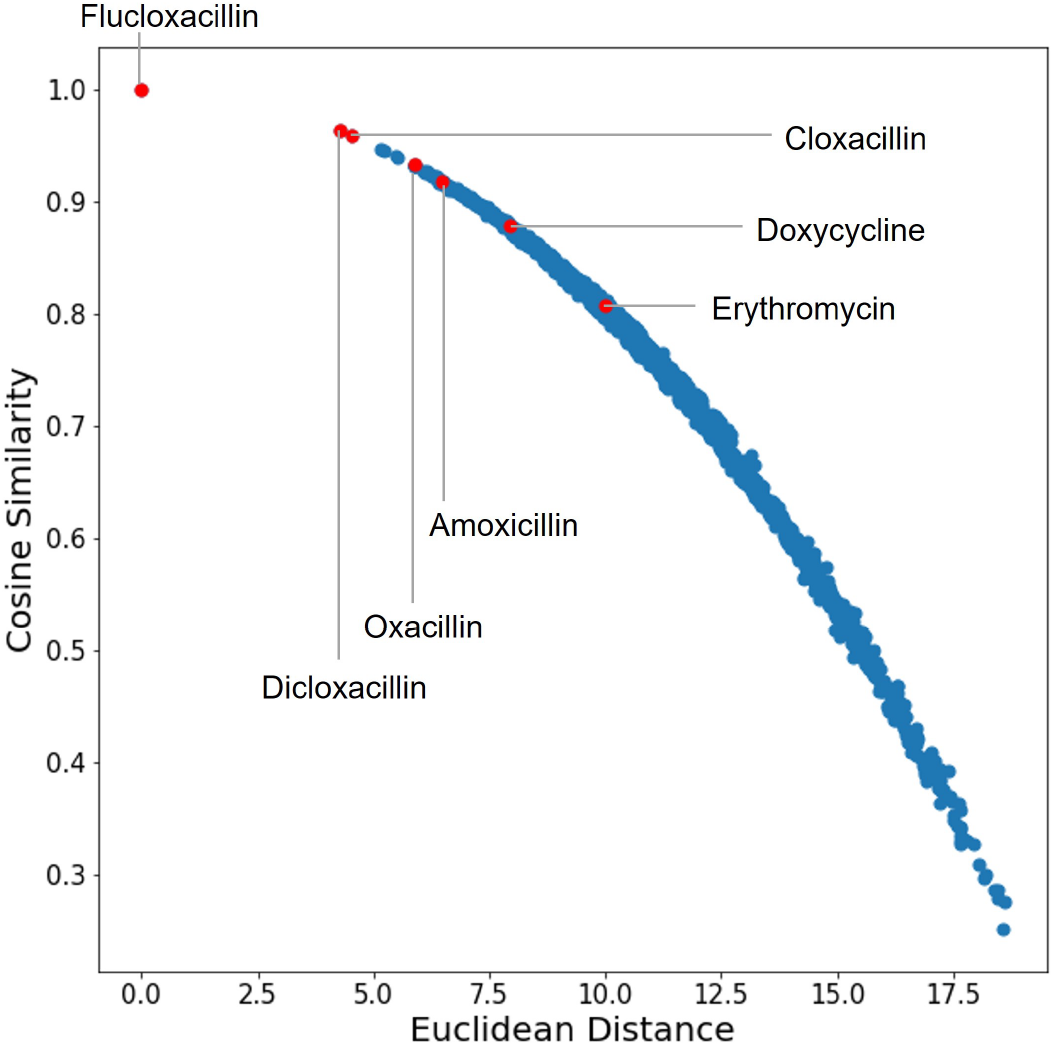
Cosine Similarity vs. Euclidean distance in the embedding space between Flucloxacillin and 1497 FDA-approved drugs.

### 3.4 Drug-drug interaction prediction

Drug-drug interaction prediction results are shown in Table 3 in terms of AUC, AUPR, precision, recall, and F-measure. In addition to our method, results of four other methods in [70] are also displayed for comparison. MM-Deacon embedding + MLP outperforms neighbor recommender method and random walk method when using substructure data only for each metric. Comparing with classifier ensemble methods that ensemble tens of neighbor recommender models, random walk models, and a matrix perturbation model on all types of similarity matrices, our method has a comparable AUC and significantly outperforms their approach for AUPR and recall metrics.

**Table 3:** Drug-drug interaction prediction metrics among 548 drugs in 5-fold cross-validation.

| Method | AUC | AUPR | Precision | Recall | F-measure |
| --- | --- | --- | --- | --- | --- |
| Neighbor recommender method using substructure data [70] | 0.936 | 0.759 | 0.617 | 0.765 | 0.683 |
| Random walk method using substructure data [70] | 0.936 | 0.758 | 0.763 | 0.616 | 0.681 |
| Classifier ensemble method (L1) [70] | **0.957** | 0.807 | 0.785 | 0.670 | 0.723 |
| Classifier ensemble method (L2) [70] | 0.956 | 0.806 | 0.783 | 0.665 | 0.719 |
| MM-Deacon embedding + MLP | 0.946 | **0.911** | **0.805** | **0.823** | **0.810** |

Since one of the aims of this drug-drug interaction dataset is to detect unknown interactions [70], we list the top 20 most potential interactions from the non-interactions predicted by our method in Table 4 and use DrugBank to verify if the false positives are true positives. As a result, 10 out of 20 are verified as true positives.

**Table 4:** Top 20 most potential interactions in the non-interaction set. True interactions verified by DrugBank are marked in **bold**.

| Rank | Drug ID 1 | Drug name 1 | Drug ID 2 | Drug name 2 |
| --- | --- | --- | --- | --- |
| 1 | DB00426 | Famciclovir | DB00741 | Hydrocortisone |
| 2 | **DB00853** | **Temozolomide** | **DB00158** | **Folic Acid** |
| 3 | DB00426 | Famciclovir | DB00327 | Hydromorphone |
| 4 | **DB00214** | **Torasemide** | **DB00468** | **Quinine** |
| 5 | DB00526 | Oxaliplatin | DB00273 | Topiramate |
| 6 | **DB00806** | **Pentoxifylline** | **DB00313** | **Valproic Acid** |
| 7 | DB00820 | Tadalafil | DB00257 | Clotrimazole |
| 8 | DB00533 | Rofecoxib | DB05271 | Rotigotine |
| 9 | **DB00199** | **Erythromycin** | **DB01142** | **Doxepin** |
| 10 | DB00455 | Loratadine | DB00612 | Bisoprolol |
| 11 | DB00346 | Alfuzosin | DB00332 | Ipratropium bromide |
| 12 | **DB00577** | **Valaciclovir** | **DB00330** | **Ethambutol** |
| 13 | DB00285 | Venlafaxine | DB01110 | Miconazole |
| 14 | **DB01204** | **Mitoxantrone** | **DB00390** | **Digoxin** |
| 15 | DB00512 | Vancomycin | DB00277 | Theophylline |
| 16 | **DB00555** | **Lamotrigine** | **DB00351** | **Megestrol acetate** |
| 17 | DB00426 | Famciclovir | DB01586 | Ursodeoxycholic acid |
| 18 | **DB00659** | **Acamprosate** | **DB00787** | **Aciclovir** |
| 19 | **DB00252** | **Phenytoin** | **DB00327** | **Hydromorphone** |
| 20 | **DB00853** | **Temozolomide** | **DB00421** | **Spironolactone** |

## 4 Discussion

In this study, we proposed a novel method called MM-Deacon for SMILES-IUPAC joint learning in a contrastive learning framework and evaluated the quality of the joint embedding space from different aspects.

The clustering of five functional groups demonstrates that the domain knowledge brought by IUPAC nomenclature has been encoded in the joint embedding space enforced by self-supervised contrastive loss. Functional groups are responsible for characteristic chemical reactions of molecules, and thus molecular representation with an awareness of underlying functional groups is beneficial for drug discovery. The clustering results of different types of embeddings in Figures 4 and 5 show that SMILES embeddings and IUPAC embeddings generated in the joint embedding space both have abilities for functional group separation. SMILES embeddings have a slightly better separation performance than IUPAC embeddings as shown in Table 1.

When searching molecules on 100K PubChem test set using cosine similarity, the embedding space trained on 10M molecules exhibits gain in performance on cross-modal search compared with training on 1M molecules as shown in Table 2. This indicates that MM-Deacon is capable of scaling to a large scale dataset, and there is still room for improvement since only 10M out of 100M of molecules from PubChem are used. Moreover, performing cross-modal molecule search in different search corpus settings (100, 10K, 100K) resulted in different recall metrics shown in Table 2. For example, for SMILES-to-IUPAC search with MM-Deacon pretrained on 10M molecules, Recall@1 on dataset of size 100 is 99.98%. This implies that when using a SMILES string of any molecule in this 100-sized dataset as a query to search across 100 IUPAC strings, there is a 99.98% chance that the IUPAC string of the same molecule with the query SMILES is the top 1 retrieved result. Likewise, Recall@1 on dataset of size 100K is 52.09%, which indicates that the chance of returning the IUPAC string of the same molecule with the query as the top 1 result is 52.09%. As the test set size goes up (from 100, to 10K, to 100K), the difficulty level of molecule search also goes up, as there are more candidate molecules in the test set. Therefore, there is a decrease of performance from 100 to 100K. Even for conducting search on 100K molecules, our model still has a Recall@5 above 85% for either SMILES-to-IUPAC or IUPAC-to-SMILES search.

From Figures 7 and 8, we can see that the top 5 retrieved molecules all have similar substructures with the query molecules, and they also appear close in the embedding space. These findings demonstrate the convergence of the pretraining in learning SMILES-IUPAC mutual information in the joint embedding space. Both SMILES and IUPAC representations can be embedded into the same joint embedding space with shared properties. Thus our model provides a choice that IUPAC strings can be used directly here instead of having to convert them into SMILES representation first for quantitative analysis. Note that unimodal search (SMILES-to-SMILES, IUPAC-to-IUPAC) is also supported in the joint embedding space, whose performance is not quantified in this study due to lack of a clearly-defined benchmark in the community [31, 34] for performance evaluation. Nevertheless, drug similarity assessment and drug-drug interaction prediction tasks show the quality of unimodal performance implicitly for SMILES representation.

In exploring the similarity relationships of drugs with Clozapine and Flucloxacillin in drug similarity assessment on FDA approved drug list, the drugs of interest in both cases all have a high rank as well as high similarity score, which also supports the claim that the underlying domain knowledge of functional groups are encoded in the embedding space. Moreover, when comparing cosine similarity with Tanimoto Similarity (RDKit) and Tanimoto Similarity (Morgan) for Clozapine, we also marked some drugs that have a high rank from cosine similarity while a low rank from Tanimoto Similarity, like Proflavine in Figure 10 and Trioxsalen in Figure 11. Like Clozapine, both Proflavine and Triosalen also have three fused rings, while their ranks in the drug list are very low when using RDKit fingerprint and Morgan fingerprint. On the contrary, cosine similarity in the embedding space has the ability to identify the mentioned structural similarities with Clozapine.

The structures of Flucloxacillin, Dicloxacillin and Cloxacillin shown in Figure 13 are nearly identical, except that Flucloxacillin has a substituent F- and a Cl- on a benzene while at the same positions, Dicloxacillin has two CI- and Cloxacillin has one Cl-. When looking at the Euclidean distances of pairs (Flucloxacillin, Dicloxacillin) and (Flucloxacillin, Cloxacillin) in the embedding space as shown in Figure 16, the distances resulted from differences in (F-, CI-) and (F-, no substituent) are both notable. This also supports that the embedding space encodes structure similarities and at the same time also has a high weight on functional groups.

Finally, when using SMILES embeddings from MM-Deacon and a simple MLP with one hidden layer for drug-drug interaction prediction task, the performance is better than neighbor recommender method and random walk method using substructure data and classifier ensemble models [70]. Moreover, similar to [70], some novel drug-drug interactions are also detected with our method. This shows that the structural information encoded in the drug embeddings can assist drug-drug interaction prediction and is superior than substructure similarity matrix obtained using structural information from PubChem.

From the evaluations on four different tasks, we have demonstrated that MM-Deacon, trained in a self-supervised manner with SMILES-IUPAC pairs, generates a molecular embedding space that fuses shared features between pairs of modalities and is a promising candidate for molecular representation.

## 5 Conclusion

In this study, we proposed a novel approach of utilizing mutual information from SMILES-IUPAC joint learning with a self-supervised contrastive loss for multimodal molecular representation learning. We evaluated our approach for molecule clustering, cross-modal molecule search, drug similarity assessment and drug-drug interaction tasks, on three publicly available datasets. Our results demonstrate that self-supervised multi-modal contrastive learning framework holds huge possibilities for chemical domain exploration and drug discovery. In future, we plan to scale MM-Deacon pretraining to larger size datasets, and also plan to investigate applicability of MM-Deacon to more downstream tasks.

